# Genome wide screen reveals a specific interaction between autosome and X that is essential for hybrid male sterility

**DOI:** 10.1101/496976

**Authors:** Yu Bi, Xiaoliang Ren, Runsheng Li, Qiutao Ding, Dongying Xie, Zhongying Zhao

## Abstract

Hybrid male progeny from interspecies cross are more prone to sterility or inviability than hybrid female progeny, and the male sterility and inviability often demonstrate a parent-of-origin asymmetry. However, the underlying mechanism of asymmetric sterility or inviability remains elusive. We previously established a genome-wide <u>h</u>ybrid <u>i</u>ncompatibility (HI) landscape between *Caenorhabditis briggsae* and *C. nigoni* by phenotyping a large collection of *C. nigoni* strains each carrying a *C. briggsae* introgression. In this study, we investigate the genetic mechanism of asymmetric sterility and inviability in both hybrid male and female progeny between the two species. Specifically, we performed reciprocal crosses between *C. briggsae* and different *C. nigoni* strains that each carries a GFP-labeled *C. briggsae* genomic fragment referred to as introgression, and scored the HI phenotypes in the F1 progeny. The aggregated introgressions cover 94.6% of the *C. briggsae* genome, including 100% of the X chromosome. Surprisingly, we observed that two *C. briggsae* X fragments that produce *C. nigoni* male sterility as an introgression rescued hybrid F1 sterility in males fathered by *C. briggsae*, indicating that at least two separate X-autosome interactions are involved in the hybrid male sterility. In addition, we identified another two *C. briggsae* genomic intervals on the Chromosome II or IV, respectively, which can rescue the inviability, but not the sterility, of hybrid F1 males fathered by *C. nigoni*, suggesting the involvement of differential epistatic interactions in the asymmetric hybrid male fertility and inviability. Importantly, backcrossing of the rescued sterile males with *C. nigoni* led to isolation of a 1.1-Mb genomic interval that specifically interacts with an X-linked introgression, which is essential for hybrid male fertility. We further identified three *C. briggsae* genomic intervals on the Chromosome I, II and III, respectively that produce inviability in all F1 progeny dependent or independent of the parent-of-origin. Taken together, we identified multiple independent interacting loci that are responsible for asymmetric hybrid male and female sterility and inviability, which provides important insights into the asymmetric HI and lays a foundation for their molecular characterization.

**Author summary:** It is common that closely related species can mate with each other, but their hybrid progeny are often sterile or inviable, especially in the male progeny. The mechanism underlying the asymmetric sterility or inviability remains poorly understood. We previously addressed this question between two nematodes, *Caenorhabditis briggsae* and *C. nigoni*, by systematic substitution of various parts of the *C. nigoni* genome with its *C. briggsae*’s equivalent followed by phenotypic examination. Here we investigate the genetic mechanism of the asymmetric sterility and inviability in the hybrid F1 male and female progeny between the two species. We achieved this through crossing a cohort of *C. nigoni* strains each carrying a substitution with *C. briggsae*, which led to differential homozygosity of the *C. briggsae* substitution in the hybrid progeny. The aggregated substitutions cover 94.6% of the *C. briggsae* genome, including 100% of the X chromosome. Surprisingly, we identified two *C. briggsae* X fragments that produced *C. nigoni* male sterility as a substitution but rescued hybrid F1 sterility in males fathered by *C. briggsae*, indicating that at least two separate X-autosome interactions are involved in the hybrid male sterility. In addition, we identified multiple genomic intervals on *C. briggsae* autosomes that can rescue the inviability, but not the sterility, of hybrid F1 males fathered by *C. nigoni*. Importantly, we isolated a 1.1-Mb genomic interval that specifically interacts with an X-linked introgression, which is essential for hybrid male fertility. We further identified three *C. briggsae* genomic intervals on the Chromosome I, II and III, respectively that produce inviability in all F1 progeny dependent or independent of the parent-of-origin. The identified interacting loci lays a foundation for their molecular characterization.

## Introduction

Postzygotic <u>h</u>ybrid <u>i</u>ncompatibility (HI) presents one of the major barriers to gene flow between species or populations, leading to reproduction isolation or its enforcement. HI commonly manifests as hybrid male sterility or overall hybrid inviability, and has undergone intensive study over in recent decades [1]. Identification of HI loci has become a major task of evolutionary biologists. Antagonistic interactions between parental alleles with differential divergence are predicted to become incompatible, hereafter termed <u>D</u>obzhansky–<u>M</u>uller <u>i</u>ncompatibility (DMI) [2,3], which is believed to be responsible for the HI phenotypes in the hybrid progeny. Genetic studies of HI have isolated various loci across phyla. A subset of the HI loci has been molecularly cloned, which had provided unprecedented insights into speciation genetics [1]. Hybrid male sterility is more common than other types of HI phenotype, which is dubbed Haldane’s rule [4]. The male sterility or viability phenotypes often depend on the parent of origin, which is called Darwin’s corollary to Haldane’s rule [5]. The two rules are well supported in the interspecies hybrids cross species [6–9]. The X Chromosome has been shown to play a disproportionately larger role in hybrid male sterility in diverse organisms, which is termed the “large-X effect” [10,11]. However, whether X-linked incompatibilities contribute disproportionately to asymmetric hybrid inviability, especially female’s inviability, has yet been established [12]. The key to understanding the genetic mechanism of these two rules is to isolate specific genomic intervals that are responsible for the asymmetric HI phenotypes in both hybrid F1 and nearly isogenic genetic backgrounds [13].

*Caenorhabditis elegans* has been intensively studied for neuron development [14], tissue differentiation [15,16] and organogenesis [17] and population genetics [18–20]. However, it has contributed little to speciation genetics, although incipient speciation seems to be obvious among various polulations [21,22]. Such study has been inhibited by the lack of a sister species that can mate with *C. elegans* and produce viable hybrid progeny [23]. An improved sampling method introduced in the early 2000s dramatically accelerated the recovery of new *Caenorhabditis* species [24–26], which led to discivery of a few new pairs of sister species that can mate and produce viable progeny [8,27]. For example, crosses between a newly identified sister species pair, *C. remanei* and *C. laten*, produced pronounced asymmetric hybrid male sterility and extensive hybrid F2 breakdown [28]. Reciprocal crosses suggested that the genetic basis of hybrid inviability is more complex than hybrid male sterility, and that hybrid male sterility in nematodes involves a single X-autosome interaction [29]. Unfortunately, mapping resolution of HI in hybrid F1 between the two species is relatively coarse.

The identification of *C. briggsae* sister species, *C. nigoni*, has paved the way for using the use this pair for speciation genetics [8]. *C. briggsae* is a close relative of *C. elegans*. They share morphology and developmental patterns [30] and are mostly hermaphrodites with occasional males, whereas *C. nigoni* is a strictly dioecious species that is mostly found in tropical areas [8]. Both species contain five autosomes and a single sex chromosome X, with XO as male and XX as female or hermaphrodite. Studies on their hybrid supported both Haldane’s rule and Darwin’s corollary to Haldane’s rule [8,9]. For example, a cross with *C. briggsae* as the father produces hybrid fertile F1 females but sterile F1 males, whereas a cross in the opposite direction produces hybrid fertile F1females only (Fig. 1). However, the genetic mechanism that underlies the asymmetric HI phenotypes remains elusive. To empower *C. briggsae* and *C. nigoni* as a model for speciation genetics, we previously generated approximately 100 visible transgenic markers that express green fluorescent protein (GFP), and each was inserted into a different part of an individual chromosome [31,32]. Because they can serve as a dominant marker for its linked *C. briggsae* genomic fragment in the hybrid progeny during backcross, these markers greatly facilitated mapping of the HI loci between the two species. Systematic backcross of all of these markers into *C. nigoni* for at least 15 generations led to a genome-wide HI landscape consisting of *C. briggsae* introgression in an otherwise *C. nigoni* background [31]. Notably, each these HI loci was identified in essentially a *C. nigoni* background. It is expected that the underlying mechanisms of HI in hybrid F1 and in the essentially isogenic genetic background that carries an introgression will differ greatly but remain poorly defined in any species.

**Fig. 1.**
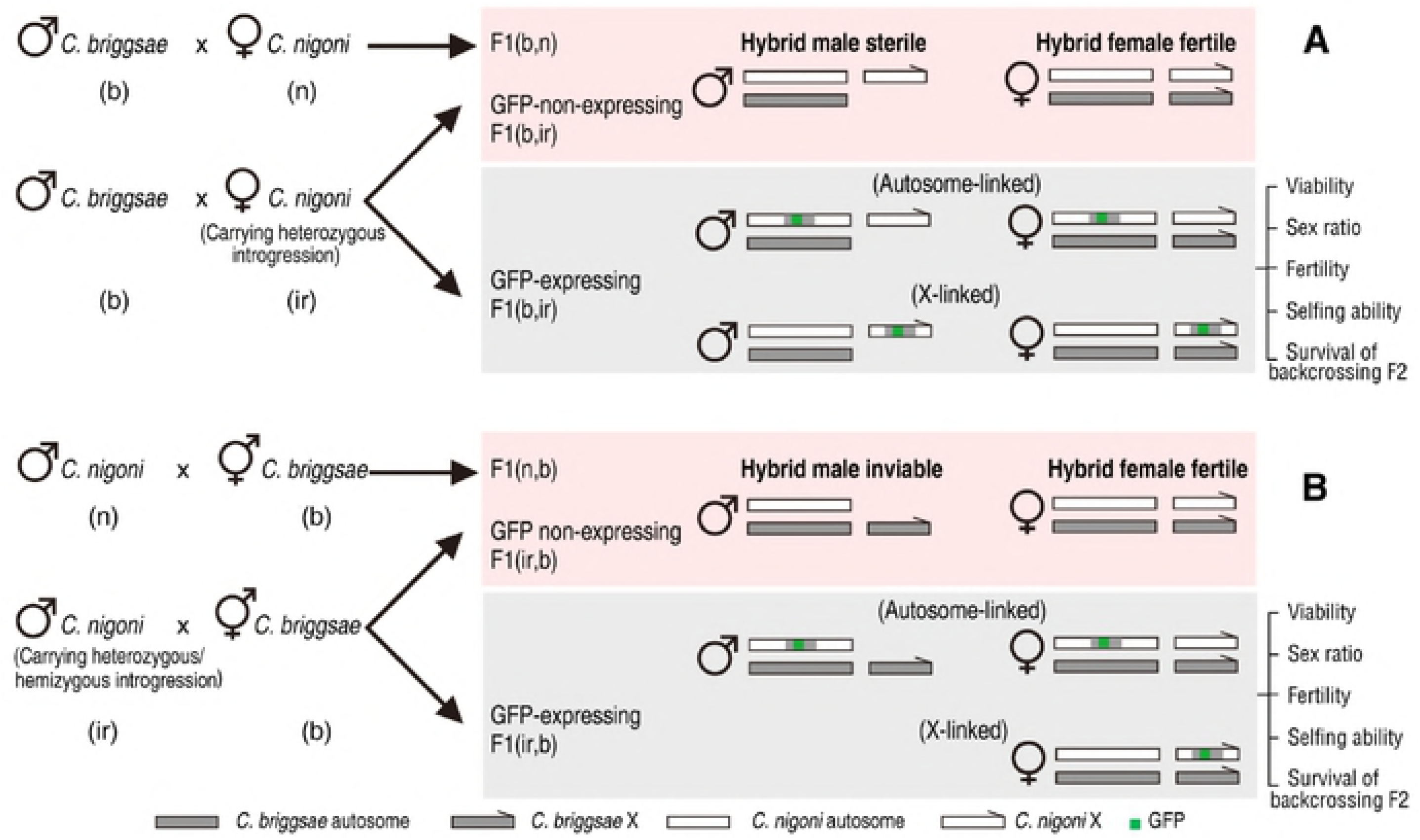
Strategy of screening for changes in hybrid incompatible (HI) phenotypes in the F1 progeny from crossing between wild-type *C. briggsae* and *C. nigoni* introgression lines as opposed to the F1 progeny from crossing between wild isolates of two parental species. Shown are schematics of crossings with *C. briggsae* (b) wild isolate (n) as a father (A) or a mother (B). Crossings with *C. nigoni* wild isolate (n) or its introgression line (ir) are shown on the top and bottom, respectively. GFP-linked introgression on autosome or X chromosome are shown separately. HI phenotypes were scored for the F1 GFP-expressing (introgression-bearing) progeny as indicated. Survival of F2 progeny in the crossing between the hybrid F1 progeny and *C. briggsae* wild isolate was also counted. Crossing progeny was named as described previously [8]. Briefly, progeny are named after its genotype in parenthesis with paternal and maternal parents listed in the left and right, respectively.

Previous study showed few involvement of mitochondia in hybrid F1 phenotype [33], indicating that most HI phenotypes result from incompatibility between nuclear genomes. A loss of function allele in *Cbr-him-8* in *C. briggsae* hermaphrodite was shown to produce viable and fertile hybrid male progeny when mated to the *C. nigoni* father [34]. Such males would allow many other genetic study, such as isolation of genes responsible for hermaphroditism. However, subsequent repeat of this cross experiment was unsuccessful [35], suggesting that the rescue of the male sterility and inviability may be an artifact from incomplete sperm depletion of *C. briggsae* mother. To search for the rescue of male sterility or inviability observed from reciprocal crosses between the two species, we took advantage of a large collection of *C. nigoni* strains each carrying an *C. briggsae*-specific introgression fragment we generated previously [31]. We performed a genome-wide screen for specific *C. briggsae* genomic fragments that could rescue the male sterility or inviability by crossing the introgression-bearing *C. nigoni* with *C. briggsae* in both directions whenever applicable. We identifed various introgression fragments that were able to rescue either the male sterility or inviability. We also identified genomic intervals that kill both hybrid males and females in cross direction-dependent or independent way. Availability of hybrid F1 fertile male further permited us to isolate a specific X-autosome interaction that is essential for hybrid male fertility.

## Results

### X-linked *C. briggsae* introgressions lead to male sterility in *C. nigoni* but rescue the male sterility in hybrid F1 progeny

One of the most common HIs between these two species is hybrid F1 male sterility between the two species with *C. briggsae* as the father (Fig. 1). We previously identified at least two independent *C. briggsae* genomic fragments from its X Chromosome that produced male sterility in *C. nigoni* as an introgression (Fig. 2A) [31]. However, the *C. nigoni* females that carry the same introgression are fertile. To systematically identify the *C. briggsae* genomic fragment that can rescue the inviability of F1(n, b) (see Materials and Methods) male, we performed reciprocal crosses between wild-type *C. briggsae* and *C. nigoni* lines that each carries an independent X-linked or autosome-linked introgression. To achieve a maximal coverage of the *C. briggsae* genome, we selected a subset of introgressions that show little or no overlap, which we generated in a previous study [31]. In addition, to narrow the interval responsible for the rescue, we included several introgressions that overlapped with those that were confirmed to produce changes in F1 phenotypes. As a result, we performed crosses for a total of 29 introgression lines, covering 94.6% of the *C. briggsae* genome, with complete coverage of the X Chromosome (Table S1).

**Fig. 2.** Substitution of specific *C. nigoni* X fragments with its *C. briggsae* counterparts produces male sterility in *C. nigoni* but rescues male sterility in F1 (b, n) hybrid. (A) Schematic representations of male fertility produced by *C. briggsae* introgressions in *C. nigoni* or by hybrid F1(b, ir). *C. briggsae* X chromosome (Cbr-X) is shown in scale as a black line. Positions for PCR primers used for genotyping are indicated as vertical line (Table S2). Blue thick bars above the X chromosome indicate previously mapped *C. briggsae* chromosome intervals responsible for the male sterility as an introgression in *C. nigoni* background. Thin bars underneath the X chromosome show the *C. briggsae* introgressions that produce hybrid F1(b, ir) fertile or sterile males, which are differentially color-coded in grey and green, respectively. For simplicity, the introgression name is shown as the 5-digits on the left without prefix “*zzyIR”*. Deduced intervals responsible for rescuing of hybrid F1 male sterility is highlighted in black boxes. (B) Quantification of hybrid F1(b, ir) male ratio in all GFP-expressing F1 progeny shown in 2A. Hybrid male fertility is color-coded as in 2A. Note that all the X-linked introgressions do not lead to a significant decrease in male ratio as opposed to the control except *zzyIR10307*, which produces a significant increase (student’s *t* test, *p* < 0.05). Number of scored animals are indicated in parenthesis. Control: male ratio in wild-type F1(b, n) hybrids. (see also Fig. S5, where all autosome-linked introgressions produce a significant decrease in male ratio). (C) Quantification of sperm size. Shown are violin plots of sperm sizes from the parental (*C. nigoni* and *C. briggsae*) males, their hybrid F1(b, n) male, the introgression-bearing F1 hybrid male between wild type *C. briggsae* father and *C. nigoni* mother carrying either of the two introgressions, i.e., *zzyIR10330* and *zzyIR10307* with fertility color-coded in dark blue (sterile) or green (fertile). Number of sperms scored are indicated. Note the size differences between wild type *C. briggsae* and *C. nigoni* males (*p <* 2.2e-16), and between the sterile and fertile hybrid males (*p <* 2.2e-16) (Wilcoxon signed-rank test). D.Quantification of sperm activation rate. Shown are bar plots of activation rate from the males in 2C. Rescued fertile hybrid males carrying an introgression demonstrate an activation rate comparable to their parental species, which is significantly higher than those of the hybrid sterile F1(b, n) males (*p <* 0.01, Two-sample student’s *t* test, unpaired). E-N. Sperm morphology and activation in parental or hybrid males. Shown are the sperms before (E - I) and after activation (J - N). From left to right: parental (*Cni* and *Cbr*) males, their hybrid F1(b, n) male, the introgression-bearing hybrid F1 male between *C. briggsae* father and *C. nigoni* mother carrying either of the two introgressions, i.e., *zzyIR10330* and *zzyIR10307*. Sperms after activation by pronase treatment are shown at the bottom. Magnified views are shown in inlet. The morphology of activation (pseudopod) seems abnormal in the sterile F1 hybrid.

To our surprise, we observed opposite phenotypes caused by either of the introgressions in *C. nigoni* and in hybrid F1 progeny; that is, a cross between *C. briggsae* males and *C. nigoni* females with the introgressions produced hybrid fertile males, but a similar cross without the introgressions produced hybrid sterile males (Figs. 1 & 2A), which suggests that an epistatic interaction between the introgression and some autosomal fragments was responsible for the observed male sterility. The gonad morphology of the rescued males was similar to that of their parental males (Fig. S1). The availability of a hybrid F1(b, n) fertile male opens the door for further genetic dissection of X-autosome interaction that is required for hybrid male fertility as described below.

In hybrid F1 adult progeny fathered by *C. briggsae*, the hybrid F1 males, which was referred to as F1(b, n) males, were not only sterile, but also demonstrated a disproportional segregation of sex ratio; that is, approximately 35% of the hybrid F1 progeny were male, and the rest were females, indicating that there are some other epistatic interactions that led to hybrid male inviability. To investigate whether the mapped genomic interval that rescued the hybrid male sterility also rescued the hybrid male inviability, we quantified the ratio of introgression-bearing males and females among all the hybrid F1 progeny from the following cross, that is, between a *C. briggsae* male and a *C. nigoni* female carrying an introgression from various parts of the *C. briggsae* X Chromosome.

We found that only one of the X-linked introgressions rescued the hybrid male inviability (Fig. 2B), suggesting that separate loci are responsible for the male sterility and inviability.

### Sperm sizes of the rescued hybrid males is more similar to that of wild type *C. nigoni* males

The sperm size of hermaphroditic *C. briggsae* is significantly smaller than that of dioecious *C. nigoni* [36], and mating between the two species with *C. nigoni* as the father sterilized the maternal *C. briggsae* [37]. We wondered which parental alleles that control sperm size are dominant. We therefore quantified the sperm size derived from the hybrid sterile, the rescued fertile males, and the parental males. We found that the sperm size of the hybrid F1(b, n) sterile males was comparable to that of *C. briggsae*, but the sperm of the rescued hybrid fertile males was significantly larger (Fig. 2C, E-I), suggesting that the *C. briggsae* alleles that control sperm size are mainly recessive. It also suggested that spermatogenesis fails during the formation of primary spermatocytes in the hybrid sterile males [36]. Sperm activation rate was comparable between two rescued males but was significantly higher than the hybrid F1(b, n) males (Fig. 2D, J-N).

### Few improvements on F2 breakdown when backcrossed to *C. briggsae*

One of prominent asymmetric HI phenotypes between *C. briggsae* and *C. nigoni* is the complete F2 breakdown of all progeny derived from the reciprocal crosses between *C. briggsae* and a hybrid F1 female (Fig. 1). The availability of hybrid F1(b, n) fertile males carrying *zzyIR10330* or *zzyIR10307* provides an opportunity to evaluate whether the F2 breakdown resulting from backcrossing to *C. briggsae* [8] still holds with an increased contribution from the *C. briggsae* genome. To test this hypothesis, we scored the brood size, survival rate, and male ratio in the progeny derived from the crosses between the rescued F1(b, ir) males that carry the X-linked introgression, *zzyIR10330* and *zzyIR10307*, and the sperm-depleted *C. briggsae* hermaphrodite (Fig. 3). About a quarter of around 50 laid F2 embryos from the cross developed into adulthood (Fig. 3 A & B). Notably, the male ratio was significantly higher than that in the F1(b, n) progeny (Fig. 3C). However, all the surviving F2 males and females became sterile with deformed gonad (Fig. S2), which suggests that the F2 inviability involved loci other than the X-linked introgressions used here.

**Fig. 3.** Quantification of fertility rescue in hybrid F1 male. Shown are bar plots of brood size (A), survival rate (B) and male ratio (C) of hybrid F2 progeny between the rescued F1 fertile males (F1 (b, 10330) and F1 (b, 10307)) and *C. briggsae* hermaphrodite or *C. nigoni* female, respectively. Number of the scored F1 males, F2 embryos and adults is indicated underneath each plot in (A), and (C), respectively. Significant deviations from control is indicated with two stars (*p <* 0.01, Student’s *t* test). #: Brood size from crossing between F1(b, n) male and *C. nigoni* female (white box)/*C. briggsae* hermaphrodite (black box). ##: Survival rate and male ratio of F1(b, n)(white box) and F1(n, b)(grey box) progeny.

We also scored the survival rate in the progeny derived from the crosses in the opposite direction for all introgressions in this study, that is, the crosses between *C. briggsae* males and the F1 progeny derived from reciprocal crossings between *C. briggsae* and *C. nigoni* carrying an autosome-or X-linked introgressions. We were intrigued to find that few progeny survived to adulthood (Fig. S3). Taken together, the increased contribution of the *C. briggsae* genome in the hybrid did not improve the fertility of the hybrid F2 progeny, thus preventing further backcross with *C. briggsae* beyond the F2 generation.

### A specific interaction between the *C. nigoni* Chromosome II and the *C. briggsae* X is responsible for hybrid male sterility in *C. nigoni*

The availability of hybrid F1 fertile male provides an opportunity to screen for specific X-autosome interactions that are responsible for the hybrid male sterility (Fig. 4A). Because the hybrid F1(b, n) males that carry either *zzyIR10330* or *zzyIR10307* are fertile, we randomly chose one of them, i.e., *zzyIR10330*, to isolate the autosomal interacting loci that are responsible for the observed rescue. We initiated the crosses by mating wild-type *C. briggsae* males and *C. nigoni* (*zzyIR10330*) females. The resulting introgression-bearing hybrid F1(b, 10330) (see Materials and Methods for nomenclature) fertile males were backcrossed with wild-type *C. nigoni* females for as many as 20 generations using the introgression-bearing fertile male. Therefore, any *C. briggsae* loci that co-segregated with *zzyIR10330* were expected to be identified by genotyping the fertile males. Given the possibility of multiple independent *C. briggsae* loci that could co-segregate with the introgression and together rescue the male sterility, we selected a single male to set up a cross for genotyping by single-worm polymerase chain reaction (PCR) using the *C. briggsae-*specific primers as described [6]. We found that the *C. briggsae* Chromosome II had specifically co-segregated with *zzyIR10330* (Fig. 4B).

**Fig. 4.** Identification of *C. briggsae* autosomal loci that interact with introgression *zzyIR10330* for rescuing the sterility of introgression-bearing *C. nigoni* male. (A) Backcrossing strategy. Hybrid fertile F1 (b, 10330) male was backcrossed with wild-type *C. nigoni* for at least 20 generations. In addition to the male fertility check, introgression bearing males were genotyped every 3-4 generations by PCR to ensure the absence of size change of the introgression, *zzyIR10330*, which could lead to the rescue of male sterility. (B) Mapping of autosomal loci co-segregating with *zzyIR10330*. Shown on the top is the genotyping results using NGS (see also Fig, Sxx), showing that the right arm of chromosome II is co-segregating with *zzyIR10330*, which is required for male fertility. Shown on the bottom is the fine mapping of interacting loci on chromosome II using existing introgressions on the chromosome II as indicated. Deduced smallest interval that rescues male sterility is highlighted in rectangle box.

Given the error-prone feature and limited resolution with the PCR-based method, it is possible that there are some other co-segregating segments that were missed by the PCR assay. To detect any other possible fragments that could co-segregate with *zzyIR10330*, we performed whole genome sequencing with Nanopore technology using the DNAs extracted from the female progeny derived from the cross between introgression-bearing males and wild-type *C. nigoni* females (see Materials and Methods). These female progeny, but not the male progeny, were expected to carry the introgression and any co-segregating loci as heterozygote in an essentially *C. nigoni* background. Consistent with the PCR-based assay, we detected only a single *C. briggsae* fragment on the right arm of Chromosome II that co-segregated with *zzyIR10330* in the hybrid rescued males (Figs. 4B and S4). The sequencing results also confirmed that the rescued male fertility did not result from a decreased length of *zzyIR10330*, but from the co-segregation of *zzyIR10230*.

The identified co-segregating *zzyIR10230* is more than a half of the size of the entire Chromosome II (Fig. 4B), which is expected to result in inviability when rendered homozygous in *C. nigoni* based on our previous introgression results [31]. To narrow the genomic interval responsible for the rescue of male sterility, we took advantage of our existing overlapping introgressions in the same region (Fig. 4B). Specifically, we set up crosses between *C. nigoni* females that carried *zzyIR10330* and six individual *C. nigoni* strains that each carried an independent introgression that overlapped with *zzyIR10230*. The presence and absence of fertile males would be indicative of the introgression’s ability or inability, respectively, to rescue the male sterility caused by *zzyIR10330* in *C. nigoni*, respectively. Our crossing results not only confirmed the significance of *zzyIR10230* in the rescue of hybrid male sterility, they also allowed us to narrow an interval of 1.1 Mb from 13.35 Mb to 14.45 Mb on the *C. briggsae* Chromosome II (cb4), which interacts with *zzyIR10330* to rescue the male sterility.

### Homozygosity of independent autosome introgressions rescue hybrid inviability of F1 (n, b) males

Another prominent HI phenotype between *C. briggsae* and *C. nigoni* is the hybrid inviability of F1(n, b) males (Fig. 1). Despite the rescue of hybrid F1(b, n) male sterility by the X-linked introgressions, these introgressions failed to rescue the inviability of the hybrid F1 (n, b) males, indicating that differential epistatic interactions are responsible for the parent-of-origin hybrid male fertility and viability.

We were able to identify two independent autosomal intervals on the right and left arms of the Chromosome II and IV, respectively, that rescued the F1(n, b) male inviability (Fig. 5A). However, the rescued males were sterile and had malformed gonad and spontaneous sperm activation (Fig. S5), which suggests that additional loci can be found elsewhere in the *C. briggsae* genome that is required for the male fertility. Contrasting the overlapping parts of the relevant introgressions allowed us to narrow the intervals responsible for the observed rescue to roughly 3.8 and 8.9 Mb on the Chromosome II and IV, respectively (Fig. 5A). To examine whether the viability rescue was complete, we scored the ratio of introgression-bearing males among all introgression-bearing progeny. We found that the ratio of the introgression-bearing males is significantly smaller than the expected 50% for all the five scored introgressions (Fig. 5B), suggesting that the rescue of viability is incomplete.

**Fig. 5.** Deviated hybrid F1 phenotypes from those of crossing progeny between parental species in the presence of autosomal introgression. (A) List of deviated phenotypes in the hybrid F1 progeny. Chromosome, introgression and its name are indicated as in Fig. 2A. Introgressions with directional and unidirectional killing of hybrid F1 progeny (both male and female) are colored in orange and red, respectively. Introgression that rescues hybrid male inviability in F1 (n, b) is colored in blue. Deduced smallest interval responsible for a given phenotype is highlighted with rectangle box. (B) Quantification of male ratio in introgression-bearing F1(ir, b) progeny with rescued male inviability (blue). Number of scored animals is indicated. Only introgressions on chromosome II and IV are found to be able to rescue the male inviability in F1 (n, b). Control: Male ratio in wild-type F1(n, b) was scored as the control group. (C) Statistics of sperm sizes in parental or hybrid males. Shown are violin plots of sperm size of wild-type males of *C. nigoni* (Cni) or *C. briggsae* (Cbr), sterile F1 hybrid (b, n) and F1(10353, b) males with the number of scored sperm indicated below. Note that the sperm size of the rescued sterile males is comparable to that of *C. briggsae* but are significantly smaller than those of *C. nigoni* male (Wilcoxon test, *p* < 2.2e-16) (Fig. 2C).

**Fig. 6.** Summary of interactions between X and autosome in determining male fertility and viability. Shown are interactions using an X-linked introgression (*zzyIR10330*) and Chr-II-linked introgression (*zzyIR10353*) as an example. The two introgressions are indicated as a GFP and RFP-linked introgression, respectively. Hybrid phenotypes are indicated. Compatible and incompatible interactions are indicated with a black and red arrow. For simplicity, only chromosome II is indicated and introgression name is only denoted by its unique identifier within its name. Note that only a single interaction is indicated, but it is possible that there are more interactions between other part of the autosome and the X-linked fragment. A. Hybrid F1(b, n) male sterility produced by an interaction between *C. nigoni* homozygous regions of *zzyIR10330* and autosome. B. Hybrid male sterility of *C. nigoni* produced by an interaction between a small interval on the *C. nigoni* chromosome II and *zzyIR10330*. C. Hybrid F1 male inviability produced by an interaction between *C. nigoni* homozygous regions of *zzyIR10353* and *C. briggsae* X. D. Hybrid F1(b, n) male inviability produced by an interaction between *zzyIR10353* and *C. nigoni* X.

To investigate the inheritance of sperm size in the rescued hybrid F1(n, b) males that carry a complete *C. briggsae* X Chromosome, we quantified the sperm size for one line of the viable males rescued by the introgression, *zzyIR10353*. Again we found that the sperm size of the rescued viable yet sterile males was comparable to that of wild-type *C. briggsae* males, but was significantly smaller than that of wild type *C. nigoni* males (Fig. 5C). These results demonstrated that a sperm is functional in the hybrid only when its size is comparable to that of wild type *C. nigoni* males. They also suggest that a part of the *C. briggsae* X other than *zzyIR10330*, is incompatible with *C. nigoni* autosomes, which is likely to explain the observed male sterility.

### Parent-of-origin dependent or independent killing of both male and female F1 progeny by different autosomal introgressions

Despite the sterility or the inviability of the hybrid F1 males, the hybrid F1 females between *C. briggsae* and *C. nigoni* were mostly viable and fertile regardless of their parent of origin (Fig. 1) [8]. Unexpectedly, we observed three independent autosomal intervals located on the Chromosome I, II, and IV, respectively, that produced inviability of both hybrid male and female progeny that carried an introgression, hereafter termed as complete inviability, when present as a homozygote in the cross progeny between *C. briggsae* males and introgression-bearing *C. nigoni* females (Fig. 5A). The intervals on the Chromosome I and II were supported by one and two introgression(s), respectively, which produced a complete inviability of F1(b, ir), whereas the interval on chromosome IV was supported by a single introgression, which produced complete inviability independent of the parent of origin. Apparently, only nuclear genomes were involved in the inviability because the viability was not improved when *C. nigoni* mitochondria were substituted by its *C. briggsae* equivalent (Table S3). Given that the three introgressions produced no inviability in the otherwise *C. briggsae* background as a heterozygote, their epistatic interactions with the remaining *C. nigoni* genome are likely to be recessive in the hybrid F1 background. No introgression on the X Chromosome produced complete inviability when present as homozygote or hemizygote, indicating that the X Chromosome is barely involved in the hybrid female viability either in hybrid F1 progeny or in the *C. nigoni* strains that carry a *C. briggsae* introgression.

To evaluate the effects of autosomal introgression on the male viability in hybrid F1 progeny, we scored the ratio of introgression-bearing males among all the hybrid F1 progeny that carried an introgression from crosses between *C. briggsae* males and *C. nigoni* females that carried an introgression. We found that the ratio of male was significantly lower than that of females in nearly all the hybrid progeny that carried an autosome-linked introgression (Fig. S6). In contrast, the ratio of males was comparable to that of females in nearly all hybrid progeny that carried an X-linked introgression (Fig. 1B), which is consistent with the observation that male-specific genes are enriched on autosome but depleted on the X Chromosome [38].

### Partial homozygosity for *C. briggsae* alleles rarely increased the incidence of selfing relative to the F1 generation

Another purpose of the crosses using the introgression strains was to explore whether the increased proportion of the *C. briggsae* genome in the hybrid can recover the selfing capability of the hybrid females. To this end, we examined the fertility of all crossing females that carried an introgression by plating a single hybrid L4 *C. briggsae* or *C. nigoni* female. We found only the L4 progeny from the cross between *C. briggsae* male and the *C. nigoni* female carrying an introgression, *zzyIR10320*, was able to produce approximately 50 eggs without further mating. A couple of the eggs hatched and grew to adulthood. However, all of them were all sterile females, indicating that increased proportion of *C. briggsae* genome in the hybrid did not recover the selfing capability.

## Discussion

Asymmetric male sterility or inviability is common in cross-species hybrids, but the mechanism responsible for these phenotypes is poorly defined from both genetic and molecular standpoints. Despite intensive study of hybrid male sterility or inviability, the mechanism of hybrid female inviability remains largely unknown. In this study, we investigated the genetic mechanism of asymmetric defects in viability and fertility of hybrid male and female progeny between hermaphroditic *C. briggsae* and dioecious *C. nigoni*.

### Differential genetic mechanism between hybrid male sterility and viability and female viability

The availability of hybrid F1(b, n) fertile males that carry a GFP-labeled introgression allowed us to dissect the genetic loci involved in hybrid F1 male sterility. By contrasting the genotypes between F1(b, n) sterile and F1(b, 10330) or F1(b, 20307) fertile males, we were able to deduce at least two independent incompatible interactions between the *C. nigoni* homologous region of *zzyIR10330* or *zzyIR10307* and the *C. briggsae* autosome in the hybrid F1(b, n) males, which are responsible for the observed sterility (Fig. 6). This is because the only difference in genotype between the two was the presence of *zzyIR10330* or *zzyIR10307* in the fertile males but not in the sterile males. This finding is consistent with the notion that the evolution of sex chromosomes is no longer dominated by the unique genetic features of the sex chromosomes themselves, but rather a result of global interactions between sex chromosomes and autosomes [39]. The study of gene birth in the *Drosophila* genome revealed disproportional translocation of X-linked genes to autosome, which are predominately testis-specific genes [40]. Intriguingly, *Drosopphila* genes translocated from autosome to the Y Chromosome are undergoing rapid degeneration and pseudogenization, suggesting a differential selection pressure of bidirectional gene translocation between the autosome and the X Chromosome. Given that the size of the hermaphroditic *C. briggsae* genome is substantially smaller than that of the dioecious *C. nigoni* and [41,42], and other *Caenorhabditis* species [43], it remains to be determined whether differential gene translocation exists between the X Chromosome and the autosomes between the two species, which is responsible for the observed X-autosome interaction.

F1(n, b) males are inviable (Fig. 1). One of the two intervals that is required for the rescue of hybrid inviability in F1(n, b) males is located on the *C. briggsae* Chromosome II, which is the same as the one responsible for the rescue of sterility in *C. nigoni* males carrying *zzyIR10330* (Figs. 2A and 5A). However, the region did not rescue the sterility of the F1(n, b) males (Fig. 5A&C), which indicates that the hybrid male fertility and viability involve different loci. The other interval required for the rescue of hybrid inviability of F1(n, b) males is located on the Chromosome IV. It is possible that the two intervals are more compatible with the *C. briggsae* X Chromosome than its *C. nigoni* syntenic parts in the hybrid F1(n, b) background, which led to the rescue.

Intriguingly, all the X-linked introgressions that were viable and fertile as homozygotes/hemizygotes in *C. nigoni* were unable to rescue the hybrid F1(b, n) male sterility, whereas those that produced hybrid male sterility as hemizygotes in a wild-type *C. nigoni* background were able to rescue the male sterility (Fig. 2A), which is consistent with the notion that multiple independent X-autosome interactions that are involved in hybrid male sterility. It should be noted that our result did not preclude a possible role of a parental effect, or a much smaller size of the X Chromosome in *C. briggsae* than in *C. nigoni*, that is responsible for the hybrid male sterility. Indeed, the parental effect was also blamed for the observed hybrid inviability between *C. remanei* and *C. laten* [29].

### Rescued fertile F1(b, ir) male facilitates precise mapping of X-autosome interaction

Precise isolation of the interacting genomic interval is challenging in the hybrid F1 animals due to the complication of the coexistence of both parental genomes. However, backcrossing of the rescued F1(b, n) males with *C. nigoni* permits isolation of the precise interacting loci responsible for hybrid male fertility in *C. nigoni* (Fig. 4A). We successfully isolated a *C. briggsae* interval of 1.1 Mb in size on the Chromosome II that interacts with *zzyIR10330*, which is essential for hybrid male fertility in an otherwise *C. nigoni* background, indicating that a specific interaction between *zzyIR10330* and the *C. nigoni* genomic interval homologous to the mapped *C. briggsae* interval that is responsible for the *C. nigoni* male sterility (Fig. 6). However, it remains possible that there could be other autosomal loci that also interact with the introgression and rescue the male sterility, but are missed in our analysis. This possibility exists because we only randomly selected introgression-bearing males from each generation for backcrossing. Some other animals could have carried different introgressions other than the one we isolated that could rescue the sterility, but they could have been missed during the backcrossing as used in this study. Given the relatively large size of the isolated genomic intervals involved in asymmetric hybrid male sterility, inviability or female inviability, it is possible that any single genomic interval may contain more than one genetic loci responsible of the observed rescue or killing. Fine mapping of these loci remains a significant challenge. Future studies should focus on the development of more reagents to allow systematic isolation of such interacting loci. One of the major limitations of our method of screening for the interacting loci is the lack of introgressions labelled with differential markers, such as GFP and red fluorescent protein (RFP). The RFP-labelled introgressions will allow direct testing of their interaction in *C. nigoni* by crossing the two *C. nigoni* strains that each carries an independent autosomal introgression labelled by RFP and *zzyIR10330*, for example, labeled by GFP. Rescue of the male sterility caused by *zzyIR10330* would be indicative of the two loci. It is worth noting that the genomic sequences between *C. briggsae* and *C. nigoni* diverge substantially from each other in both sequence identity and size [41,44], which significantly inhibits DNA recombination frequency. Selective loss of male-specific genes in hermaphroditic species and its subsequent reinforcement further complicates the precise mapping of HI loci using backcrossing. Therefore, development of overlapping introgressions in different colors provides an alternative to narrow the genomic interval for a given phenotype (Fig. 4A).

### Possible role of incompatibility between piRNA pathways in non-directional killing of hybrid F1 progeny

Although Haldane’s rule and Darwin’s corollary to Haldane’s rule are well supported in the hybrids between *C. briggsae* and *C. nigoni*, the hybrid females are fertile regardless of the crossing direction (Fig. 1). We were surprised to find that in the hybrid F1 progeny that is homozygous for the *zzyIR10018* region, both male and female progeny are inviable regardless of the parent of origin (Fig. 5A). Notably, the introgression spans the entire region of the second piRNA cluster on the *C. briggsae* Chromosome IV [45,46], which raises the possibility that incompatibility of these piRNAs between *C. briggsae* and *C. nigoni* that is responsible for the hybrid F1 inviability caused by crossing in both directions. Consistent with this notion is another piRNA cluster unique to *C. briggsae* on the Chromosome I [46]. The homozygosity of the region covering this cluster led to inviability in the hybrid F1 male and female progeny regardless of the parent of origin (Fig. 5A). The exact mechanism of directional killing remains to be determined.

## Materials and Methods

### Nematode strains and maintenance

The following strains were used in the study: *C. briggsae* (AF16), *C. nigoni* (JU1421), *C. nigoni* cytoplasm-replaced hybrids (ZZY10357) and 29 *C. nigoni* strains carrying a GFP-flagged *C. briggsae* introgression (Table S1). ZZY10357 carrying a complete set of *C. nigoni* chromosomes and *C. briggsae* cytoplasm were generated by backcrossing *C. briggsae* hermaphrodites with *C. nigoni* male worms for 20 generations. Therefore, only mitochondria from *C. briggsae* but not from *C. nigoni* are present in *C. nigoni* background as judged by genotyping with *C. briggsae*-specific PCR primers. All strains were maintained at a constant temperature of 25°C o n 1.5% agar NGM (Nematode Growth Medium) plates seeded with the *E. coli* strain OP50.

### Crossing setup and nomenclature of hybrid

The paternal and maternal strains were named as described previously [8]. Specifically, the genotype of the hybrid F1 progeny was indicated by their parent of origin in the parenthesis after “F1”, which were separated by a comma. The letter “b” and “n” were used to denote *C. briggsae* and *C. nigoni*, respectively. The word “ir” was short for introgression and a 5-digit number derived from its corresponding strain name was used to denote a specific strain. For, example, F1(b, n) represented F1 hybrids from crossing between *C. briggsae* male and *C. nigoni* female animals, while F1(n, b) represented F1 hybrids from the opposite cross direction. When introgression-bearing *C. nigoni* male/female animals were crossed with *C. briggsae* hermaphrodite/male animals, their hybrids were referred to as F1(ir, b) and F1(b, ir), respectively with “ir” denoting “introgression”. For a specific introgression, a unique number derived from its name (Table S1) was used to replace “ir”. For example, F1(b, 10330) denoted hybrid F1 progeny derived from the cross between *C. briggsae* male and *C. nigoni* female carrying an introgression *zzyIR10330.*

### Viability

Hybrid viability was measured as the number of hybrids could that survive to adulthood. Specifically, F1(b, ir) or F1(ir, b) hybrid viability was measured by scoring the number of GFP-expressing and non-GFP-expressing male and female adults, which was used to evaluate the effect of an introgression on the hybrid viability relative to that of hybrid F1(b, n) or F1(n, b). Note that hybrid inviability was exclusively found in F1(n, b) male hybrids but not in F1(b, n) male or female hybrids from both cross directions.

### Male ratio

Given there was some intraspecific variations between male ratio scored using different strains [9], male ratio was scored in the hybrids derived from cross between wild isolates of *C. briggsae* (AF16) and *C. nigoni* (JU1421) in both cross directions as the control. For an introgression of interest, the hybrid male ratio was scored as the proportion of hybrid male adult expressing GFP out of all GFP-expressing hybrid progeny. The final ratio was calculated from the average of at least 3 replicates, each being scored for at least 150 animals.

### Fertility

Hybrid fertility was defined as the ability to produce embryos after mating with the opposite sex from either *C. briggsae* or *C. nigoni*. Note that hybrid sterility was exclusively found in F1(b, n) male but not in F1(b, n) or F1(n, b) female (Fig. 1) [8]. F1(n, b) male was unsuitable for fertility test due to hybrid inviability except in the case of rescued inviability. Fertility of hybrid males bearing an introgression of interest was evaluated by mating GFP-expressing males with *C. nigoni* L4 females or *C. briggsae* sperm-depleted hermaphrodites for 24 hours. Fertility of hybrid females bearing an introgression of interest was identified by mating GFP-expressing L4 females with *C. briggsae* or *C. nigoni* males for 24 hours. Fertility was quantified as brood size, which as calculated from the average number of embryos produced by at least three individual worms in their whole lifespan.

### Screen for selfing ability

*C. nigoni* carrying an introgression (GFP-expressing) were mated with *briggsae* for 12 hours in both directions. Parents were killed. Laid eggs were allowed to L4 stage followed by transferring individual females into separate plates. Number of embryos and grown animals were counted after 72 hours cultivation on NGM plate with food. The F2 animals were mated *C. briggsae* and *C. nigoni* for fertility.

### Survival rate of backcrossing F2

In cross between F1 female hybrids and *C. briggsae* males, hundreds of backcrossing F2 embryos were laid but less than 1% of them were able to develop to adulthood. The crossing was set up by mating GFP-expressing F1(b, ir) or F1(ir, b) L4 females with *C. briggsae* males for 24 hours, followed by transferring the F1 female hybrids every 12 hours till they stopped laying more egg. The number of embryos and survived adults was counted. The survival rate was calculated as average ratio of F2 adults out of the total F2 embryos in 3 replicates, with at least 150 embryos counted in each replicate.

### Microscopy

A Leica LSM 510 confocal laser microscope was used for taking all DIC micrographs. Worms were paralyzed in M9 buffer with 0.05M sodium azide. Spermatids from dissected worms were incubated in sperm medium (SM) containing 50 mM HEPES pH7.8, 50 mM NaCl, 25 mM KCl, 5mM CaCl2, 1 mM MgSO4, 1 mg/ml BSA) [47].

### Sperm size quantification

GFP-expressing well-fed male was picked under stereo microscope. Spermatids were obtained by dissecting male worms in SM followed by imaging with the Leica microscope. ImageJ [48] was used to automatically measure cross-sectional area of spermatids using the function “particle analysis”. For each experiment, sperm size was measured for 3 to 4 males, each producing 85 to 648 spermatids.

### Sperm activation and its quantification

Sperms were activated in freshly made SM containing diluted pronase (Sigma-Aldrich) in a final concentration of 200 ng/μl. Micrographs were taken within 5-30 minutes after dissection. Activation rate of wild-type *C. briggsae* or *C. nigoni* male sperms was scored as control. Sperm activation rate was calculated as percentage of average activation rate (number of activated spermatids divided by number of all sperms in a microscopic view) of 3 to 4 male adults.

### Nanopore sequencing for genotyping zzyIR10230 that rescued the male sterility of ZZY10330

Fertile GFP-expressing F20 male hybrids were mated with L4 stage *C. nigoni* female worms in three batches. Genomic DNA was extracted from the females of 21^st^ generation backcrossing progeny. A total of 371 F21 adult females were picked for DNA extraction and Nanopore pore sequencing as described [41]. GFP-expressing F20 male hybrids were genotyped with PCR to ensure the boundary of X-linked introgression was the same with *zzyIR10330*, which produced the male sterility in *C. nigoni* background. It was expected that 100% of the F21 female worms carried the GFP-flagged *zzyIR10330* as a result of sex-linked segregation of the introgression, and 50% carried the non-GFP-flagged *zzyIR10230* both as a heterozygote. The Nanopore sequencing data were submitted to NCBI under BioProject number of PRJNA507071.

## Acknowledgments

We thank Chung Wai Shing and Cindy Tan for logistic support and members of Zhao’s lab for helpful comments. Some strains were provided by the CGC, which is funded by NIH Office of Research Infrastructure Programs (P40 OD010440).

## Supporting information captions

Fig S1. Morphology of males with various genetic background.

Fig S2. Morphology of hybrid F2 progeny derived from crossing between the rescued hybrid F1 fertile male with *C. nigoni* female or *C. briggsae* hermaphrodite.

Fig S3. Quantification of survival rate of crossing F2 progeny between the *C. briggsae* male and the female of F1 (b, ir) or F1 (ir, b).

Fig S4. Determining the boundaries of co-segregating fragment that rescues the male sterility produced by zzyIR10330 in *C. nigoni* using NGS.

Fig S5. Morphology of hybrid F1 sterile males from crossing between introgression-bearing *C. nigoni* male and *C. briggsae* hermaphrodite. Note that F1 (n, b) males are inviable.

Fig S6. Male ratio in introgression-bearing F1(b, ir) hybrids.

Table S1. Details of the introgression strains used in hybrid F1 phenotypic screening.

Table S2. List of the newly designed genotyping primers based on *C. briggsae* genome assembly (cb4) in addition to those listed previously (Bi, et al., 2015, PLoS Genetics)

Table S3. Statistics of hybrid progeny resulting from crossings that produced complete inviability.

